# Fractionation of sex differences in human cortical anatomy

**DOI:** 10.1101/2025.09.10.675377

**Authors:** Hyo M. Lee, Siyuan Liu, Elisa Guma, Elizabeth Levitis, Rebecca Shafee, Gabrielle Dugan, François M. Lalonde, Liv Clasen, Alex DeCasien, M. Mallar Chakravarty, Jason P. Lerch, Konrad Wagstyl, Angela Delaney, Armin Raznahan

## Abstract

Humans show reproducible sex differences in regional cortical volume (CV), but it remains unclear how these arise from underlying sex-biases in the two biologically dissociable determinants of CV: surface area (SA) and cortical thickness (CT). Moreover, limited access to experimental methods in humans has hindered direct studies of the causal drivers of regional sex differences in the human cortex, although rodent models have argued for both chromosomal and gonadal contributions to sex-biased mammalian cortical development. Here, we first use structural neuroimaging data in two independent human cohorts (combined N=1,754; 967 females) to quantify and spatially resolve the differential contributions of SA and CT to observed sex differences in CV. These dissociable facets of sex-biased cortical organization are highly reproducible and align with distinct functional networks and histo-molecular signatures. We then leverage complementary neuroimaging data in clinical case-control cohorts (combined N=313) featuring variations in X and Y chromosome dosage (sex chromosome aneuploidies) and testicular hormone production (isolated GnRH deficiency) to establish that regions of sex-biased CV, SA and CT in humans are enriched for congruent anatomical effects of X-chromosome dosage (e.g., primary sensory and insular cortices) and gonadal hormones (e.g. dorsomedial frontal and temporo-parietal-occipital regions). Taken together, these findings substantially advance both the breadth and granularity of our understanding regarding sex-biased cortical organization in humans – disambiguating sex effects on regional CV, SA and CT and nominating their potential genetic and endocrine causes.

## Introduction

Sex* modulates the prevalence and presentation of diverse brain-related health conditions^1^. For example, childhood-onset conditions like autism and attention deficit hyperactivity disorder show male-biased prevalence^2,3^, whereas mood and anxiety disorders develop a female-biased prevalence in adolescence^4,5^. Such sex differences likely stem from a complex and interactive mix of biological, psychological and social influences on human brain organization. To date, the bulk of available research on sex differences in human brain organization has come from *in vivo* neuroimaging^6–11^, which represents our primary means of studying the living human brain at scale.

The majority of neuroimaging studies on sex differences in human brain organization have considered anatomical properties measured by *in vivo* structural magnetic resonance imaging (sMRI). Although findings vary^12^, in part due to diverse methodological factors^13,14^, there is now convergent evidence from best-practice computational analyses in large independent datasets that females and males show statistically significant and reproducible mean differences in regional cortical volume (CV). These CV differences are of small-to-medium effect sizes and exist above and beyond the well-known sex differences in overall brain size^14^. Alongside this consolidation of evidence for sex differences in CV, there has been continued accumulation of evidence that the two determinants of CV - surface area (SA) and cortical thickness (CT) - have highly divergent evolutionary^15–17^, genetic^18–20^, environmental^21^, developmental^22^ and cellular^23,24^ determinants. For example, imaging genetic studies have found low genetic correlation between SA and CT with genes influencing SA showing enrichments in prenatal neural progenitor cells and those influencing CT showing enrichments for mid-fetal myelination, branching or pruning^21^. This profound dissociability of SA and CT means that analyses of CV alone can conflate two very different sets of biological processes and obscure isolated changes in SA or CT^25^. To date, however, the most literature on sex differences in human cortical anatomy has focused on CV, with only a handful of studies including parallel analysis of CV, SA and CT^6,9,25,26^, and none directly inter-relating these patterns of sex effects in a quantitative manner. As a result, we do not know how sex effects on regional CV, SA and CT vary in their spatial distribution, effect size, reproducibility or alignment with independent measures of cortical structure and function. Addressing these questions would substantially refine our understanding of sex-biased human brain organization and – given the highly divergent neurodevelopmental underpinnings of SA and CT – provide a more biologically grounded starting point for probing causal models.

Our current causal models for sex-biased mammalian brain development are largely based on experimental studies in rodents (especially mice), with a longstanding emphasis on gonadal sources of sex-biased brain development, and more recently incorporated consideration of sex chromosome dosage influences^27^. Relevant for mechanistic models for sex-biased cortical anatomy in humans, recent sMRI work in the Four Core Genotype (FCG) mouse model - in which the sex chromosome complements (XX vs. XY) are decoupled from gonadal types (testes vs. ovaries) - has indicated that both of these factors can influence regional brain volume including regions of sex-biased CV^28–30^. Although the lack of access to experimental approaches in humans has hindered formal tests of the causal roles of these two biological factors in shaping sex-biased brain anatomy, humans display certain rare medical conditions that could potentially enable causal testing for sex chromosome dosage and gonadal influences on brain organization^31^. In particular, conditions like sex chromosome aneuploidy (SCA)^32–34^ and isolated gonadotropin-releasing hormone (GnRH) deficiency (IGD)^35^ can model phenotypic effects of sex chromosome dosage variation and hypogonadism, respectively, in humans. To date, however, no studies have combined the study of SCAs and IGD with mapping of normative sex differences to determine if regions of sex-biased anatomy show congruent anatomical sensitivity to variations in sex chromosome dosage and gonadal hormones. Such a study design could help to prioritize candidate mechanistic models for sex-biased human brain development in a regionally resolved manner – especially if conducted in a manner that disambiguates biologically dissociable aspects of brain anatomy like SA and CT.

Here, motivated by these considerations, we use computational analyses of cortical anatomy by sMRI in two large cohorts of typically developing controls (combined N=1,754) as well as case-control clinical cohorts of SCA and IGD (combined N=313) to: (i) systematically parse reproducible normative sex differences in regional CV, SA and CT; (ii) show that normative sex-biases in CV can be fractionated onto regionally varying contributions from sex differences in SA and CT that are concentrated within cortical systems with distinct postmortem molecular and *in vivo* functional signatures; and (iii) establish that sex differences in cortical anatomy, especially SA and CT, are preferentially colocalized with congruent neuroanatomical effects of X chromosome dosage and gonadal hormones.

Taken together, these efforts provide a more phenotypically fine-grained understanding of sex differences in human cortical anatomy and nominate specific functional and molecular features that may differentially predict the distinct regional patterns of anatomical sex difference observed for SA and CT. Moreover, by establishing the existence of a spatial coherence between anatomical sex differences and anatomical effects of sex chromosome dosage variations and gonadal hormones, our study lends support to the hypothesis that sex chromosomes and hormones shape regional sex differences in human brain anatomy – as they have been experimentally proven to do in mice^29^.

## Results

### Sex Differences in Regional Cortical Volume Reflect Spatially Dissociable Alterations of Cortical Surface Area and Thickness

We used the Human Connectome Project (HCP) Young Adult S1200 release as our primary sMRI dataset for surface-based mapping of sex differences in cortical anatomy (N=1,085 aged 22-37 years, 592 female, **Sup Table 1**). All scans were processed through the FreeSurfer pipeline to measure CV, SA and CT for each of 360 regions (**Methods**) from the Glasser multimodal cortical parcellation^36^. For each measure, we then applied linear models to quality-controlled outputs from this pipeline to estimate the effect size and statistical significance of sex effects on regional cortical measures after controlling for age, Euler index (an index of cortical surface reconstruction quality), and global anatomical phenotypes (total CV, total SA and mean CT) (**Methods**). We observed spatially graded sex differences in CV, SA and CT (**Figure 1A**) above and beyond sex differences in total CV, total SA and mean CT, respectively (**Methods, Sup Figure 1**). The spatial distribution of these sex differences was strongly correlated across the cortical sheet between CV and SA, moderately correlated between CV and CT, and weakly correlated between SA and CT (**Figure 1B**, R=0.84, R=0.48 and R=0.2, respectively). The spatial correlations between sex differences in CV vs. SA and CV vs. CT both survived two complementary permutation-based tests for statistical significance – one based on recomputing sex difference maps and their inter-correlations after 10,000 permutations of sex (p_PERM_, **Methods**, **Figure 1B**), and another based on a Spin Test^37,38^ involving 10,000 random spatial rotations of maps for regional sex differences (p_SPIN_, **Methods**, **Figure 1B**) – both of which survived control for multiple comparisons across the 3 inter-map comparisons using Bonferroni correction (p_PERM-BF_ and p_SPIN-BF_). Thus, considering the cortical sheet as a whole, sex differences in regional CV appear to be more strongly driven by sex differences in regional SA than CT, and there is little similarity between the topography of sex-biased SA and CT.

**Figure 1.**
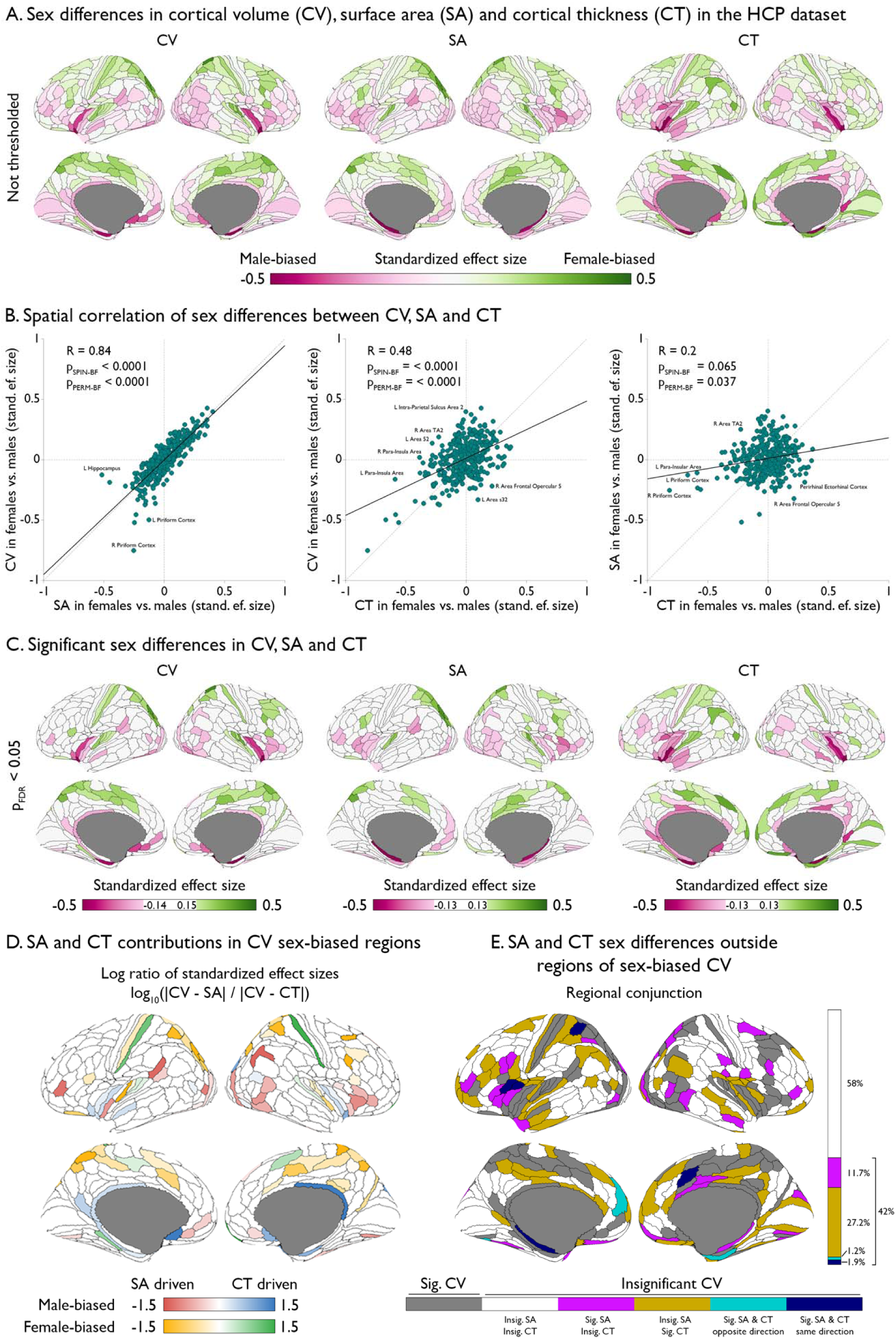
Sex differences in cortical volume, area and thickness (CV, SA, CT) in the HCP dataset. **A**. Standardized effect sizes across 360 cortical regions showing sex differences in CV, SA and CT after controlling for covariates (age, Euler index, and global CV, SA and CT, respectively). **B**. Scatter plots showing cortex-wide spatial correlations for the standardized effect sizes in sex differences between the 3 cortical features. The vertical and horizontal dotted lines are positioned where x and y axes are zero to indicate the four quadrants. The dotted diagonal denotes the y=x identity line. The solid line indicates the y∼x least squares regression fit, with the legends showing Pearson correlation (R), and Bonferroni-corrected statistical significances (across the 3 plots) of the empirical p values from 2 permutation tests (p_PERM-BF_ sex-shuffling; p_SPIN-BF_ spatial map rotation). **C**. Standardized effect sizes surviving false discovery rate (FDR) correction at p_FDR_<0.05 across 360 cortical regions showing statistically significant sex differences in the 3 cortical features. **D**. A log ratio of the absolute effect size disparity between sex differences in CV and SA and between sex differences in CV and CT quantifies the relative contribution of SA and CT to each region of statistically significant sex differences in CV. Separate color bars encode these relative contributions (i.e. SA driven vs. CT driven) for regions of female- and male-biased CV. **E**. Conjunction map showing whether sex differences in SA and CT are significant in the cortical regions lacking a statistically significant sex difference in CV. The bar inset indicates the relative proportion of cortical regions outside the sex-biased CV that show each possible combination of statistical (in)significance for sex differences in SA and CT.

Sex differences in regional cortical anatomy surviving false discovery rate (FDR) correction^39^ for multiple comparisons across cortical regions at p_FDR_<0.05 involved substantial proportions of the cortical sheet for all 3 phenotypes (28.6% for CV, 28.3% for SA and 33.1% for CT; **Figure 1C, Sup Table 2**) and ranged in absolute effect size (|d|) as follows: 0.14-0.75 for CV, 0.13-0.52 for SA and 0.13-0.82 for CT. Across all 3 phenotypes, female-biased regions (i.e. those with a statistically significantly larger mean value in females) tended to localize to dorsal cortical regions and male-biased ones to ventral regions – but we observed both convergences and dissociations between the 3 phenotypes within this broad dorsoventral pattern. All 3 phenotypes were female-biased in bilateral superior parietal and dorsal prefrontal regions as well as primary auditory cortex bilaterally, and male-biased within bilateral insular and entorhinal cortex. CV and SA were both female-biased across the right lateral frontal cortex and male-biased in bilateral temporo-occipital and left temporo-parieto-occipital junctions as well as the right frontal operculum. CV and CT were both female-biased in bilateral primary sensory cortex and male-biased in bilateral retrosplenial cortex. These observations suggested regional variation in the relative balance of SA and CT contributions to CV sex differences – motivating us to directly quantify the relative contributions of SA and CT to observed sex differences in regional CV. As a preparatory step for this quantification, we first established that the spatial profile of observed sex differences in CV is indeed a direct product of those observed for SA and CT (**Sup Figure 2**) –verifying the geometric expectation that group differences in SA and CT together determine group differences in CV. We then devised a continuous log ratio score (**Methods**) that is positive when CV sex differences are primarily driven by CT and negative when they are primarily driven by SA (**Methods**). By calculating this ratio for all cortical regions showing statistically significant sex differences in CV (**Figure 1D**), we found that: female-biased CV is driven by CT in the primary sensory cortex and primary, secondary and tertiary auditory cortices, but by SA in parietal and medial frontal cortices (e.g., area 24d); and male-biased CV is primarily CT-driven in limbic cortices (hippocampus, posterior cingulate, entorhinal cortex and area 25), but primarily SA-driven in posterior inferior parietal cortex, posterior 47r area, perisylvian language area, anterior ventral insular area, fundus of the superior temporal gyrus, temporo-parieto-occipital junction.

Separate estimation of CV, SA and CT also enabled us to quantify the extent of sex-biased cortical anatomy that is “missed” by solely focusing on the more commonly researched measure of CV. Specifically, well over a third (42%) of those cortical regions lacking a statistically significant sex difference in CV showed a statistically significant sex difference in SA, CT or both these features (**Figure 1E**). This observation further highlights the value of considering all three primary geometric properties of the cortical sheet – its volume, area and thickness - in parallel.

### Regional sex differences in CV, SA and CT show good within-sample reproducibility in a sample size dependent manner

We quantitatively assessed the within-sample reproducibility of sex differences in CV, SA and CT using 10,000 instances of split-half, sex-balanced random subsampling of the HCP dataset at varying sample sizes and without replacement (**Figure 2A**). In each instance, sex differences were recomputed in each independent split-half of the HCP dataset and the resulting cortical sex difference maps were correlated between halves. For all 3 features, the mean cortex-wide spatial correlation of sex differences between independent split-halves steadily strengthened with increasing sample size - asymptotically reaching a plateau towards the maximum possible HCP split-half sample size of 450. At this split-half sample size, we observed moderate to strong reproducibility of these sex differences in human cortical organization - reflected in cortex-wide correlations of sex differences as follows: CV mean ± s.d. R=0.67 ± 0.04; SA R=0.62 ± 0.05; CT R=0.75 ± 0.02. The variance in split-half correlations was notably lower for CT than CV and SA – a disparity which strengthened at increasing sample sizes. Moreover, all 3 phenotypes showed substantially weaker and more variable spatial correlations between split-halves at lower sample sizes – with some observed correlations for CV and SA still approaching zero at split-half sample sizes up to 200. This observed sample size dependence of reproducibility for sex differences in mean regional CV, SA and CT (**Figure 2A**) show that these sex differences are a reliable property of the cortical sheet that is recovered with ever greater stability as data increases. They also help to plan future study designs and make sense of the observation that reported regional sex differences in cortical anatomy can be highly variable at smaller sample sizes even when other methodological features are held constant^12^.

**Figure 2.**
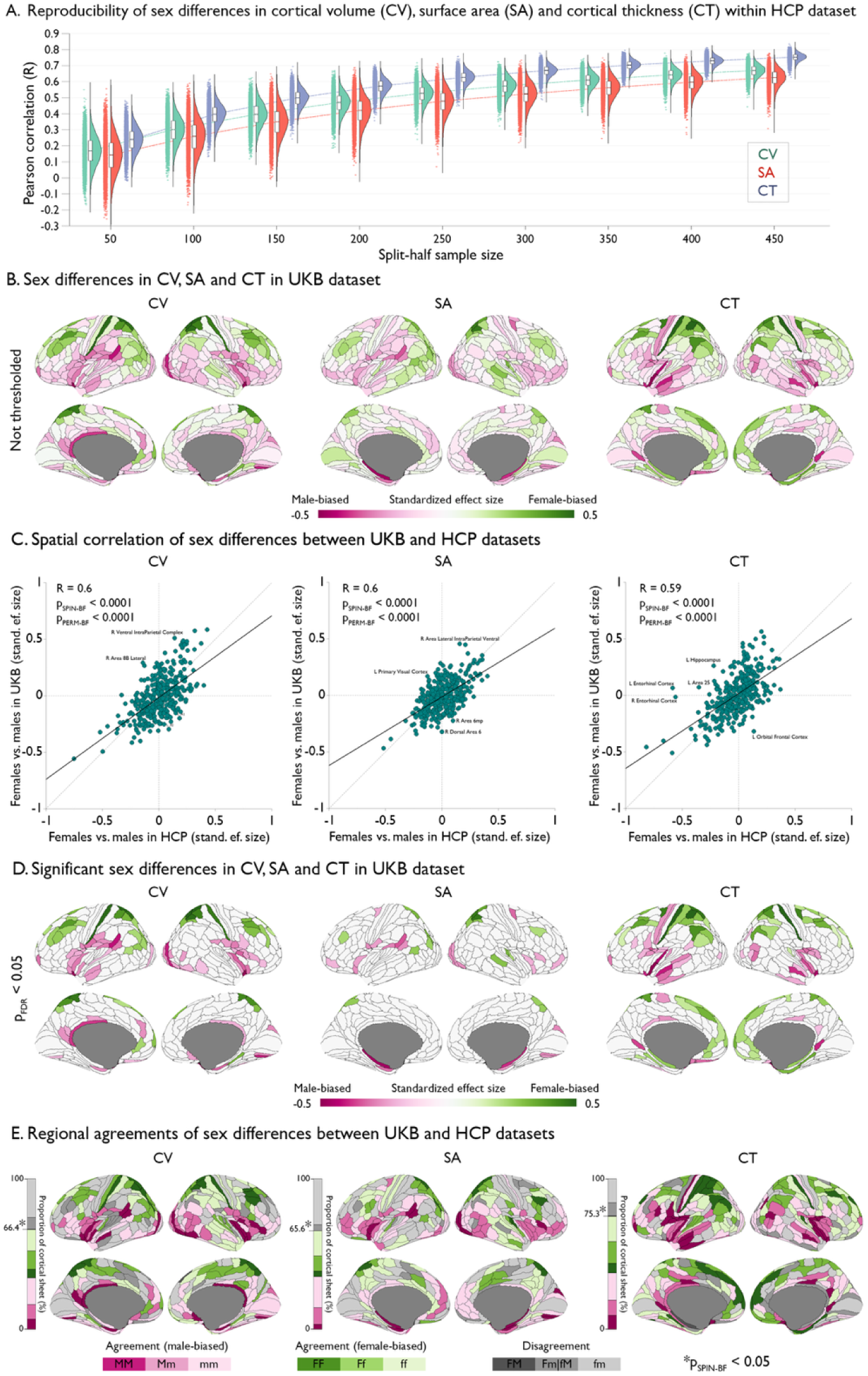
Reproducibility of sex differences in cortical volume (CV), surface area (SA) and thickness (CT) within and across datasets. **A**. Cortex-wide spatial correlations (R) between sex differences in CV (green), SA (red) and CT (blue) from 10,000 instances of split-half, sex-balanced random subsampling without replacement of varying sample sizes from 50 to 450 with increments of 50 showing reproducibility within the HCP dataset. In each instance, sex differences were recomputed in each independent split-half and the resulting cortical sex difference maps were correlated between halves. **B**. Standardized effect sizes across 360 cortical regions showing sex differences in CV, SA and CT after controlling for covariates (age, Euler index and global CV, SA and CT) in the UKB dataset. **C**. Scatter plots showing cortex-wide spatial correlations for the standardized effect sizes in sex differences between the 3 cortical features. The vertical and horizontal dotted lines are positioned where x and y axes are zero to indicate the four quadrants. The diagonal dotted line indicates the identity between the two cortical features. The solid line indicates the regression line, with the legends showing Pearson correlation (R), Bonferroni-corrected statistical significances at 0.05 relative to spin-based spatial (p_SPIN-BF_) and sex permutations (p_PERM-BF_) (**Methods**). **D**. Standardized effect sizes thresholded at p_FDR_<0.05 across 360 cortical regions showing statistically significant sex differences in each of these 3 cortical features within the UKB dataset. **E**. Conjunction maps showing 3 levels of agreement between statistically significant anatomical sex differences in the UKB and HCP datasets for CV, SA and CT. Green and pink regions are concordantly female- and male-biased between datasets, respectively (dark: both datasets significant in the same direction; neutral: either significant in the same direction; light: both insignificant in the same direction). Gray regions are discordantly sex-biased between datasets (dark: statistically significant and in opposite directions between datasets; neutral: opposite directions between datasets and statistically significant in one; light: opposite directions between datasets, but not statistically significant in either). Bars show the proportion of cortical regions falling within each of these conjunction categories. Asterisks indicate Bonferroni-corrected significance at p_SPIN-BF_<0.05 for the observed proportion of concordant cortical regions between HCP and UKB using a spin-based spatial permutation test (**Methods**).

### Regional sex differences in CV, SA and CT show good between-sample reproducibility

We quantified the out-of-sample reproducibility of those sex differences in cortical anatomy observed in the HCP sample (**Figure 1**) by re-estimating sex differences through the same image processing and computational pipeline (**Methods**) in an independent dataset gathered using different MRI machines in a different country (UK Biobank, UKB^40^, n=669, 375 females, from the 5-year age band of 44-50 years closest to HCP, **Sup Table 1**). The effect size maps for regional sex differences in mean CV, SA and CT in UKB (**Figure 2B**) showed remarkable qualitative similarity to those in the HCP (**Figure 1A**), which was quantitatively confirmed by observing a Pearson R∼0.6 correlation in regional sex differences for each phenotype between HCP and UKB datasets (**Figure 2C**). These between-dataset correlations in the topography of sex differences in all 3 measures of cortical anatomy survived testing for Bonferroni-corrected statistical significance using both sex- and spin-based spatial permutations (p_PERM-BF_ and p_SPIN-BF_, respectively; inset values **Figure 2C**; **Methods**). As per analyses in the HCP dataset, we applied FDR correction for multiple comparisons across cortical regions to localize regions of statistically significant (p_FDR_<0.05) sex differences in CV, SA and CT within the UKB dataset (**Figure 2D**). This revealed a qualitatively highly similar spatial distribution of regions showing statistically significant sex-biases in CV, SA and CT in the UKB dataset as had been observed in the HCP dataset (**Figure 1C**). We formalized this comparison by generating a conjunction map of sex differences in UKB and HCP for each of the 3 features (**Figure 2E**). These conjunction maps defined 3 levels of agreement between the two datasets (UKB and HCP both showing a statistically significant sex difference in the same direction; UKB and HCP both showing the same direction of sex difference, but only reaching statistical significance in one dataset; UKB and HCP both showing the same direction of statistically non-significant sex difference) and 3 levels of disagreement (UKB and HCP showing a statistically significant sex difference in opposite directions; UKB and HCP showing a sex difference in opposite directions which reached statistical significance in one dataset; UKB and HCP showing statistically non-significant sex differences in opposite directions). All 3 features showed agreement between datasets in over 65% of cortical regions (66% of regions for CV, 66% for SA and 75% for CT), a proportion which was statistically significantly elevated based on spin-based spatial permutation test (sex-based permutation being inapplicable due to varying numbers of statistically significant regional sex-biases across permutations, **Methods**). Moreover, of those remaining cortical regions showing a disagreement between datasets, most (CV: 76%, SA: 88%, CT: 64%) involved statistically non-significant sex differences in both datasets. The sole regional measure (i.e. 0.09% of all regional measures queried) showing an opposite direction of statistically significant sex-biased volume in HCP and UKB datasets (female>male and male>female, respectively) was left hippocampal volume – a notable exception given evidence for sex-biased age effects on this structure^8^ and the 35.6 year disparity in mean age between HCP and UKB cohorts. Thus, sex differences in regional CV, SA and CT not only show good within sample reproducibility (**Figure 2A**) but are also largely stable between independent cohorts scanned at different ages, in different countries, and on different MRI machines (**Figure 2B-E**).

### Regions of sex-biased SA and CT possess specific functional and molecular signatures

Given the spatial dissociability of sex differences in regional SA and CT (**Figure 1B**) and their regionally heterogenous contributions to regional sex differences in CV (**Figure 1D**), we next tested if observed regional sex differences in SA and CT also diverged in their functional and molecular signatures – using spin-based spatial comparisons (**Methods**) between maps of sex-biased cortical anatomy in the primary HCP dataset (**Figure 1A**) and independent annotations of cortical functional connectivity networks (Yeo-Krienen 17 network parcellation^41^) and cortical gene expression [derived from Allen Human Brain Atlas (AHBA)^42^ and mapped to the cortical surface^43^, **Methods**].

The mean standardized effect size of sex on cortical anatomy was greater than expected by chance (nominal p_SPIN_<0.05, **Figure 3A bar plots, Methods**) for a total of 6 distinct functional connectivity networks derived from resting state functional MRI data (**Figure 3A left inset**). Three of these overlaps survived correction for multiple comparisons across the 3 cortical features (p_SPIN-BF_<0.05). At this strictest level of statistical significance, the specific network overlaps varied as a function of the direction of anatomical sex-bias involved. We observed significantly male-biased anatomy within the limbic network for CV and SA and the sensory/ventral attention network for CT, alongside significantly female-biased anatomy within the dorsal attention network for all three anatomical phenotypes and within the sensorimotor network for CT. Thus, sex differences in cortical anatomy partly cohere with the topography of cortical resting state networks, and observed network enrichments vary depending on both the anatomical phenotype and direction of sex-bias being considered.

**Figure 3.**
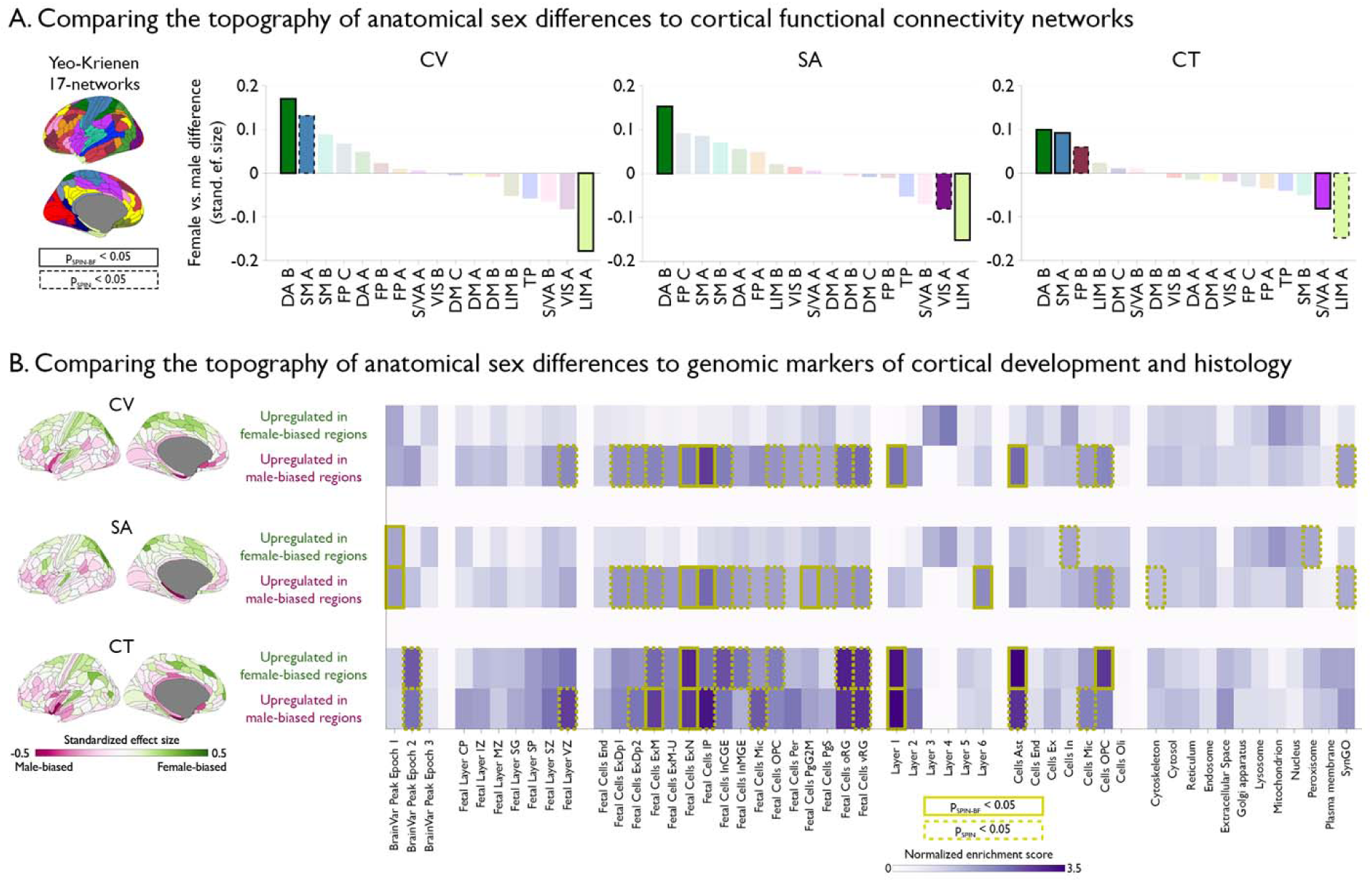
Sex differences in cortical anatomy show varying enrichments for diverse functional networks and molecular signatures depending on their direction and whether they involve cortical volume (CV), surface area (SA) or thickness (CT). **A**. The mean standardized effect sizes of sex differences in each of the 17 Yeo-Krienen resting-state functional networks (left cortical map) were compared to spin-based spatial permutations of sex difference and functional network maps to determine statistical significance. Networks that survived Bonferroni correction across 3 cortical features (p_SPIN-BF_<0.05) are shown in solid boxes, and those that were nominally significant (p_SPIN_<0.05) are shown in dotted boxes. The cortical map is color coded by the regional mapping of the functional networks. SM: somatomotor, DA: dorsal attention, FP: frontoparietal, S/VA: salience/ventral attention/ LIM: limbic, DM: default mode, VIS: visual, TP: temporoparietal. **B**. Gene Set Enrichment Analysis^44^ tests for the enrichments of the Allen Human Brain Atlas genes^43^ that correlate with the magnitude of female- and male-biased in CV, SA and CT. Statistical significance is again shown at two thresholds (nominal p_SPIN_<0.05 and Bonferroni-corrected p_SPIN-BF_<0.05) as derived from spin-based spatial permutation tests (**Methods**). The cortical maps show standardized effect sizes of sex differences in CV, SA and CT from Figure 1A as a reference. CP: cortical plate, IZ: intermediate zone, MZ: marginal zone, SG: supragranular, SP: subplate, VZ: ventricular zone, End: endothelial, ExDp: excitatory deep layer, ExM: excitatory maturing, ExN: excitatory migrating, IP: intermediate progenitor, InCGE: interneuron caudal ganglionic eminence, InMGE: interneuron medial ganglionic eminence, Mic: microglia, OPC: oligodendrocyte progenitor cell, Per: pericytes, PgG2M: cycling progenitor (G2- and M-phase), PgS: cycling progenitor (S-phase), oRG: outer radial glia, vRG: ventricular radial glia. Ast: astrocyte, Ex: excitatory, In: inhibitory, Oli: oligodendrocyte, SynGO: synaptic gene ontologies and annotations.

To characterize the molecular signatures for regions of sex-biased anatomy and annotate these at different levels of cortical organization, we used Spin Tests to rank all measured genes in the AHBA dataset of postmortem gene expression in the adult human brain^42^ by the correlation of their regional cortical expression^43^ with each direction of sex-biased anatomy for each of the 3 cortical phenotypes considered (yielding 6 ranked gene lists, one for each of: female-biased CV, male-biased CV, female-biased SA, male-biased SA, female-biased CT, and male-biased CT) (**Methods**). We used rank-based Gene Set Enrichment Analyses (GSEA^44^) to test for enrichment of highly ranked genes for a previously collated library of 51gene sets tagging different epochs of brain development, cortical layers, cell types and subcellular compartments^43^. The p value for these enrichment tests in GSEA was compared to that for repeated enrichment analyses using 10,000 random spin-based spatial permutations of each map of sex-biased cortical anatomy to derive an empirical p_SPIN_ value, which was then subjected to additional Bonferroni correction for multiple comparisons across the 6 ranked gene lists for each annotational gene set to derive an empirical p_SPIN-BF_ value.

This analytic pipeline revealed that sex-biases in SA and CT track with distinct genomic signatures of cortical development and histology (**Figure 3B**) – with observed signatures for SA sex differences being most similar to those for CV (in keeping with the stronger spatial correlation between regional sex differences in SA and CV, **Figure 1B**). Regions of sex-biased CT - regardless of the direction of this sex difference – were characterized by heightened expression of genes tagging Layer 1 of the adult cortex and prenatal post-mitotic neurons. Regions of male-biased CT showed additional enrichments for genes tagging prenatal mitotic neurons while regions of female-biased CT showed additional enrichments for expression markers of adult astrocytes and oligodendrocyte precursor cells. In contrast, regions of both female and male-biased SA in the adult cortex were enriched for genes showing peak expression in early prenatal brain development (8-24 post-conception week) – a signature not seen for regions of sex-biased CT. Regions of male-biased SA in the adult cortex were additionally enriched for genes tagging cycling prenatal progenitor cells, mitotic and post-mitotic prenatal neurons and postnatal cortical Layer 6. These multiscale annotations use intrinsic gene expression profiles to achieve a “virtual histology”^43,45^ of sex-biased cortical anatomy – specifying expression markers of distinct molecular and cellular features that spatially covary with sex differences in CV, SA and CT. Although these observations are based only on spatial correlations and therefore unable to test causal hypothesis, they provide one rational and quantifiable basis for prioritizing cellular and molecular features that are colocalized with – and therefore well placed to shape – sex differences in cortical morphology.

### Regions of sex-biased human cortical anatomy show congruent sensitivity to variations in X-chromosome dosage and testicular hormone production

The analyses above fractionate sex differences in cortical anatomy by: (i) separately mapping sex-biases in CV, SA and CT; (ii) establishing the within- and between-sample reproducibility of these sex-biases; and (iii) indicating that they tag cortical regions with distinct functional and molecular signatures. We next sought to build on this fractionation by probing potential causal mechanisms for sex differences in these dissociable aspects of cortical anatomy. Specifically, we compared maps of sex-biased CV, SA and CT with complementary maps of how these same cortical features are impacted by variations in sex chromosome dosage and hypothalamic-pituitary-gonadal axis function.

To model sex chromosome dosage effects on human cortical anatomy we harnessed neuroimaging data in males with XXY and XYY syndromes as compared to matched euploidic XY controls (Total N = 272; 99 XXY males and 92 XY controls; 34 XYY males and 47 XY controls, **Sup Table 1)**. These conditions occur in 1-2^46,47^ and 1^48^ in every thousand live male births, respectively, and are well-known to impact regional CV, SA and CT above and beyond their effects on overall brain size^32,33,49^. To model gonadal influences on human cortical anatomy, we harnessed a previously unpublished neuroimaging dataset in males with isolated GnRH deficiency (IGD, estimated prevalence 1 in 8,000 to 30,000 boys^50,51^) and healthy male controls (N=41, 19 IGD, **Sup Table 1**). The contrast between control males and those with IGD captures the influence of intact testicular hormone production on cortical anatomy because IGD is characterized by hypogonadism due to a congenital deficit in hypothalamic drive for pituitary release of the gonadotrophins that are required to drive typical testicular development and function^52^. Compared to controls – and not withstanding heterogeneity in postnatal steroid treatment – at the group level, males with IGD have been exposed to reduced levels of circulating testicular steroid hormones during those perinatal periods which are thought to be especially important for testosterone-dependent masculinization of the mammalian brain^51^. This period of hypogonadism is extended further for those males with IGD who are not diagnosed and treated with steroid replacement until later childhood and adolescence^53^.

Using identical imaging analysis procedure as those used to map sex differences in the HCP dataset (**Methods**), including control for the corresponding global phenotypes, we mapped the statistically significant effects of increasing X chromosome dose (i.e. XXY vs. XY, termed “X”, p_FDR_<0.05, **Figure 4A**), increasing Y chromosome dose (XYY vs. XY, termed “Y”, p_FDR_<0.05, **Figure 4B**) and intact testicular hormone production (controls vs. IGD, termed “T”, nominal p<0.05 as no regions showed differences surviving FDR correction, **Figure 4C**) on regional cortical CV, SA, and CT, as well as their non-thresholded effects (**Sup Figure 3**). To assess potential sex chromosomal and gonadal contributions to normative sex differences, we computed cortex-wide Pearson correlation of X, Y and T effects with sex-biases. This showed significant correlations of sex-biases with X (R=0.44, R=0.31 and R=0.25 for CV, SA and CT), Y (R=0.29 for CV) and T (R=-0.27 and R=-0.27 for CV and SA). To provide a spatially resolved view, we then color-coded all regions of statistically significant sex differences in CV, SA and CT (i.e. those regions colored in **Figure 1C**) for their congruence with the regional effects of X, Y and T (**Figure 4A-C, lower panels**). Here, we defined congruent effects as those where the direction of observed X, Y and T effects is compatible with these effects causing the observed normative sex difference (i.e. where female-biased: XXY>XY, XYY<XY or control<IGD; where male-biased: XXY<XY, XYY>XY or control>IGD). In these analyses, we differentiated between “hard” congruences (where observed X, Y and T effects reached statistical significance, top rows **Figure 4 A,B,C**) and “soft’ congruences (where observed X, Y and T effects fell below statistical significance, **Sup Figure 3 A,B,C**). This conjunction analyses revealed significant proportions of the cortical sheet where X effects are congruent with sex-biased CV, SA and CT (74.8%, 67.7% and 63% for CV, SA and CT; p_SPIN-BF_<0.05 for all), with hard congruence found in insula, temporo-parietal-occipital region, superior temporal visual area, primary sensory cortex, medial frontal and dorsal parietal region. Y effects are congruent in insignificant proportion of the cortical sheet with sex-biased CV, SA and CT (35%, 39.2% and 51.3% for CV, SA and CT) with only a few hard congruences found in e.g. parieto-occipital sulcus and area p32. T effects are congruent in a significant proportion of cortical sheet with sex-biased CV and SA (71.8% and 74.5% for CV and SA; p_SPIN-BF_<0.05), with hard congruences sparsely distributed across retrosplenial complex, temporo-parietal-occipital region, medial frontal and dorsal parietal regions. Hence, regional influences of X dosage are aligned with both sex-biased SA and CT, while those of testicular hormone production are preferentially aligned with sex-biased SA.

**Figure 4.**
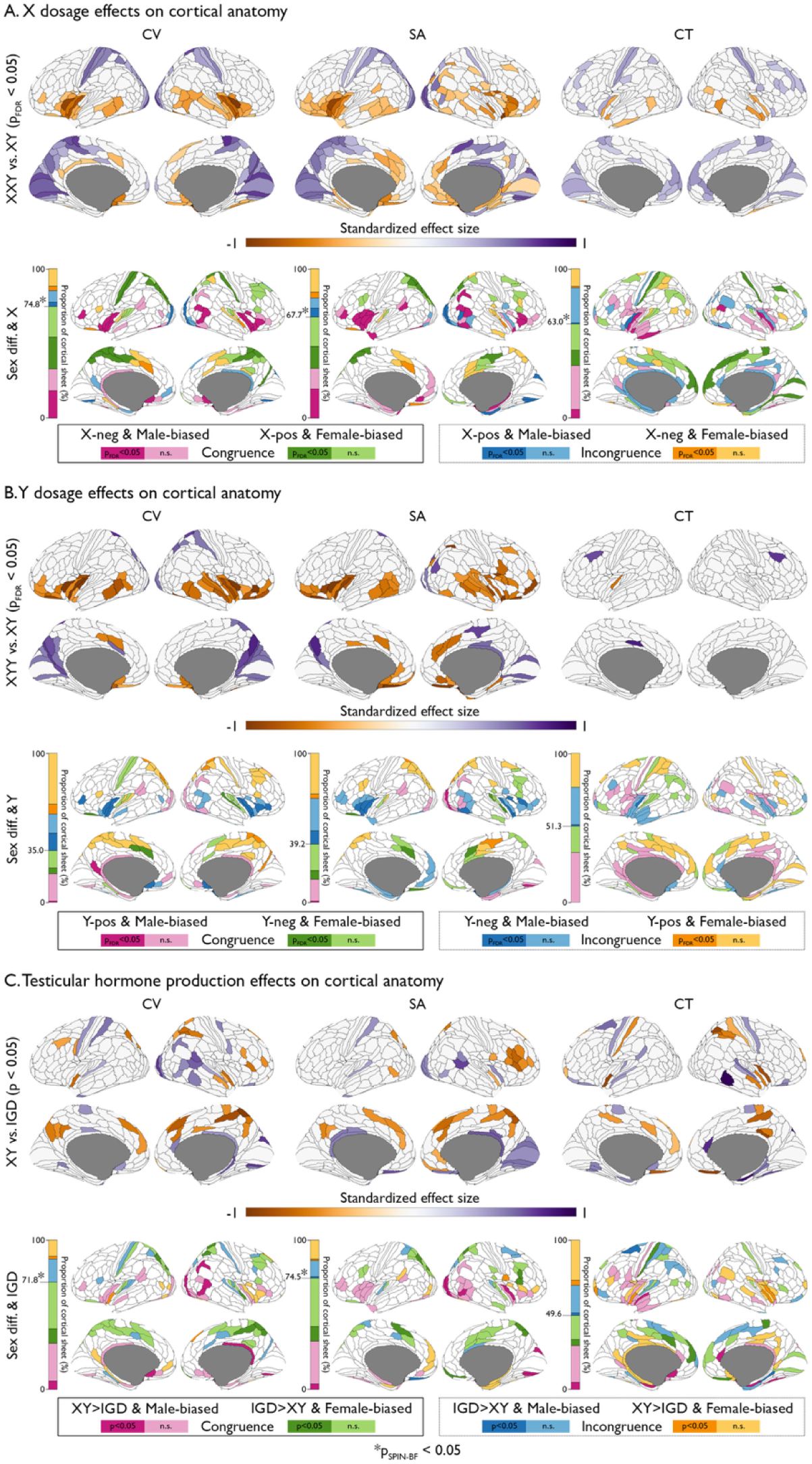
Effects of sex chromosome dosage and testicular hormone production on regional cortical anatomy. **A**. First row: standardized effect sizes across 360 cortical regions showing statistically significant (p_FDR_<0.05) X dosage effects (i.e. XXY vs. XY differences) on CV, SA and CT after controlling for age, Euler and global phenotypes (total CV, total SA and total mean CT). Second row: region-wise congruence between the effects of sex differences and X dosage. Congruence is defined when the X dosage effect is significantly in congruent direction (darker) or insignificantly in congruent direction (lighter) for female-(green) and male-biased (pink) regions. Incongruence is defined as the effect showing significantly incongruent direction (darker) and insignificantly incongruent direction (lighter) for female-(orange) and male-biased (blue) regions. The bars show the proportions at which these conjunction cases make up the cortical sheet – providing numerical insets for those proportions which are statistically significantly larger than expected under the null hypothesis. Asterisks indicate Bonferroni-corrected significance at 0.05 relative to spin-based spatial permutations of the HCP sex difference maps. **B**. Same as panel **A** but for statistically significant (p_FDR_<0.05) Y-chromosome dosage effects (i.e. XYY vs. XY differences). **C**. Same as panel **A** but for statistically significant (p<0.05) effects of intact testicular hormone production (i.e. differences between male controls and males with IGD).

Together, these results lend support to a theoretical causal model where regional sex differences in human cortical anatomy are contributed to by both sex chromosome and gonadal differences between males and females. Moreover, our findings specify subsets of sex-biased cortical regions where each of these effects may be operating – helping to refine causal hypothesis.

## Discussion

Our study advances understanding of sex-biased cortical organization in humans along four key directions as detailed below.

First, we systematically disambiguate sex effects on SA and CT – the two distinct dimensions of cortical anatomy that together determine the most studied aspect of cortical anatomy: its volume. Statistically significant sex effects on regional CV, SA and CT include medium-to-large effect sizes and exists above and beyond the robust sex effects on global CV, SA and CT. Considering the cortical sheet as a whole, we find evidence for spatially uncoupled sex effects on SA and CT, with sex effects on CV being more linked to those on SA than CT (in keeping with the dominance of SA vs. CT as a driver of interindividual CV variation^54^. Despite this general dominance of SA as a driver of CV sex differences, a region-by-region analysis reveals several instances of CV sex differences that appear to be entirely driven by CT, including primary sensory cortex and primary, secondary and tertiary auditory cortices (female-biased) as well as paralimbic cortices (male-biased). Given that SA and CT reflect largely distinct genetic^18–20^, cellular^23,24^ and developmental^22^ processes, our regional fractionation of CV sex differences according to the relative contribution of SA and CT provides a form of mechanistic stratification of sex effects of cortical anatomy. We also show how separate analysis of CV, SA and CT identifies many more statistically significantly sex-biased cortical regions than would be detected by analysis of CV alone (59% of cortical regions sex-biased considering all 3 features vs. 29% considering CV alone) – underlining the importance of combining information across different imaging derived phenotypes.

Second, we comprehensively chart the moderate-to-strong intra- and inter-sample reproducibility of sex-biased CV, SA and CT. By systematically varying sample sizes in our within-sample reproducibility analyses, we help to put prior reports of inconsistent neuroimaging sex differences^12^ into clearer context: even for morphometric sex differences as reproducible and substantial in effect size as those we find for CV, SA and CT, the discovered topography of sex differences can be still highly unstable at sex-balanced sample sizes of 400 individuals. The presented sample size dependence helps to guide study design and interpretation of future work.

Third, having disambiguated sex differences in regional CV, SA and CT, we show that these different aspects of sex-biased cortical organization flag cortical systems with distinct functional and histo-molecular signatures. It is important to stress that observing spatial alignment between a morphometric sex difference and another facet of cortical organization – such as the resting state functional connectivity or gene expression studied herein – does not establish any causal relationships. Interpretation of such spatial alignment is further complicated by the fact that the datasets from which we draw functional and transcriptomic cortical annotations are distinct from those in which we map neuroanatomical sex differences. Nevertheless, we reason that those cortical features which show close spatial covariation with anatomical sex differences are more likely to be causally related to the anatomical sex-biases than those that do not. Given this logic, it is notable that those resting-state functional connectivity networks, which show enrichment for anatomical sex-biases (sensorimotor, dorsal and ventral attention and limbic), have also been reported to show sex differences in their spatial distribution^55^ and connectivity^56^ and have been implicated in the neurobiology of multiple sex-biased neuropsychiatric disorders^57–59^. These observations help to target the design of those future studies that will now be needed to comprehensively characterize regional inter-relationships between sex differences in brain and behavior. Analogously, our transcriptomic annotation of sex differences in regional CV, SA and CT (albeit using gradients of gene expression in a predominantly adult male dataset^42^) draw attention to specific cell types and cortical layers – potentially helping to tailor the challenging future work that will be required to resolve the proximal histological and cellular determinants of sex-biased cortical anatomy as measured by MRI.

Finally, as an initial probe for potential causal drivers of sex-biased cortical anatomy, we combine our systematical mapping of regional sex differences in CV, SA and CT with studies of these same cortical features in rare clinical cohorts that capture sex chromosome dosage and gonadal effects on brain anatomy. Neuroimaging of sex chromosome aneuploidies (SCAs) and isolated GnRH deficiency (IGD) provides important causal information regarding the influences of X-chromosome dosage, Y-chromosome dosage and testicular hormone production on human cortical organization – offering a means of inquiring whether regions of sex-biased CV, SA and CT also show anatomical sensitivity to these potential drivers of phenotypic sex differences. It is important to note, however, that: effects of sex chromosome dosage variation in aneuploidy may not recapitulate those that operate in the euploidic context between XX and XY individuals; and, while IGD is a powerful model for gonadal insufficiency, it is also complicated by phenotypic variations in severity of hormone deficiency, heterogeneous hormonal replacement regimens and the potential for primary alterations in neurodevelopment^60^. Nevertheless, considering the cortical sheet as a whole, we observe a statistically surprising tendency for regions of sex-biased anatomy to show congruent effects of varying X-chromosome dosage and hypothalamic-pituitary-gonadal axis function.– consistent with the hypothesis that these sources of sex-biased biology may shape neuroanatomical sex differences in humans^61^ as they have been experimentally shown to do in rodents^62^. This observation opens a door to region-by-region annotation of sex differences for the presence of plausibly causal effects of sex chromosome and gonadal effects (i.e. those that are in a congruent direction with the observed sex-difference). Specifically, a subset of regional neuroanatomical sex differences also shows statistically significant and directionally congruent effects of X-chromosome dosage, including the primary sensory cortex (female-biased CV and CT), intraparietal sulcus (female-biased CV and SA), bilateral insula (male-biased CV, SA and CT), and angular gyrus (male-biased SA). Under the same logic, we find evidence for contributions of testicular hormone production to a subset of neuroanatomical sex differences including those in bilateral intraparietal cortex (female-biased CV and SA), temporo-parieto-occipital junction (male-biased CV and SA), and rostromedial prefrontal cortex (male-biased CV and CT). Taken together, these finding add to growing evidence for potential X-chromosome and gonadal influences on sex-biased human brain anatomy^11,63^, and help to inform follow up studies probing expression of sex chromosome genes and sex steroid pathways in specific regions of sex-biased brain anatomy. Although the present study has focused on chromosomal and gonadal contributions to normative anatomical sex differences, it is also important to consider potential contributions from those many experiential factors that differ in their prevalence and form between females and males (including gender-related variables) and carry the capacity to shape brain development^64^. Moreover, sex differences in brain anatomy are likely to vary over development^7,8,10,22,65–68^, as are the potential causal roles for chromosomal, gonadal and experiential factors^69,70^ – making it important to revisit the efforts initiated here in longitudinal datasets where available.

Notwithstanding the aforementioned caveats and limitations, our study provides an unprecedentedly comprehensive characterization and phenotypic fractionation of sex differences in human cortical anatomy and combines multiscale annotations and parallel studies in rare clinical groups to nominate candidate causal mechanisms for these sex differences. Our findings emphasize the importance of disambiguating different facets of cortical anatomy in the mapping and mechanistic annotation of sex differences.

## Methods

### Participants

Normative sex differences were mapped in the primary HCP dataset and out-of-sample reproducibility of these differences was assessed in a subset of the UK Biobank (UKB) dataset. The HCP sMRI dataset was downloaded from the HCP 1200 release (see ^71^ for details of HCP recruitment procedures) and included 1,085 healthy participants (592 females, 22-36y; 493 males, 22-37y)). The UKB sMRI dataset included 669 participants (375 females, 45-49y; 294 males, 46-49y) within the 5-year age range (44-50y) closest to that of the HCP sample (see www.ukbiobank.ac.uk and ^40^ for details of UKB recruitment procedures).

Sex chromosome dosage effects on cortical anatomy were mapped using an sMRI dataset including 272 participants from two age-matched (two sample t-test p > 0.05) case-control sex chromosome aneuploidy (SCA) cohorts: XXY (n = 99, 6-25y; 92 control participants, 5-24y); and XYY (n = 34, 7-25y; 47 control participants, 5-24y). All individuals with SCA were reconfirmed to show a non-mosaic XXY or XYY karyotype by metaphase spread (or inspection of existing community records evidencing this in the 10 carriers unable to give blood). Participants were recruited through parent support organizations, the National Institute of Mental Health website and the National Institutes of Health (NIH) Healthy Volunteers office. All participants had normal radiological findings and no history of brain injuries. All control participants were screened using a structured interview to ensure no history of neurodevelopmental or psychiatric disorders. All participants gave consent (or assent with parental consent as appropriate), and all protocols were completed at the NIH Clinical Center in Bethesda, Maryland.

The IGD dataset includes 41 participants from an age-matched case-control cohort (19 male IGD patients, 14-41y; 22 age-matched control patients, 13-42y). They were recruited through referrals from collaborators and local physicians or self-referrals with advertisement of the study on the NIH and *Eunice Kennedy Shriver* National Institute of Child Health and Human Development websites as well as ClinicalTrials.gov. Participants had a confirmed diagnosis of IGD with failure to undergo normal, age-appropriate pubertal development and low testosterone in the setting of low or inappropriately normal luteinizing hormone and follicle-stimulating hormone levels. Potential participants with additional pituitary hormone deficiencies or taking medications known to cause acquired hypogonadotropic hypogonadism (e.g. corticosteroids, chronic opioids) were excluded.

The demographic characteristics of HCP, UKB, SCA and IGD datasets are detailed in **Supplemental Table 1**. The analyses of the datasets and the research protocol were approved by the National Institute of Mental Health Combined Neuroscience Institutional Review Board.

### MRI Acquisition

T1-weighted (T1w) 3D magnetization-prepared rapid acquisition gradient-echo (MPRAGE) scans were acquired for all participants. The HCP participants were scanned using 3T ConnectomeScanner (adapted from Siemens Skyra, Siemens Healthineer) with the following parameters: 0.7 mm isotropic resolution, FOV = 224 mm, TR = 2400 ms, TE = 2.14 ms, TI = 1000 ms, FA = 8°, bandwidth = 210 Hz per pixel, and GRAPPA = 2 (see^72^ for details). The UKB participants were scanned using 3T Siemens Skyra with the following parameters: 1 mm isotropic resolution, FOV = 256 mm, TR = 2000 ms, TE = 2.01 ms, TI = 880 ms, FA = 8°, 240 Hz per pixel, echo spacing = 6.1 ms and GRAPPA = 2 (see^73^ for details). The SCA participants were scanned using 3T MR750 scanner (General Electric) with the following parameters: 1 mm isotropic resolution, FOV = 256 mm, TR = 2530 ms, TE = 3.5ms, TI = 1100 ms, FA = 7°, 195.3 Hz per pixel and GRAPPA = 2. The IGD participants were scanned using 3T MR750 scanner (General Electric) with the following parameters: 1 mm isotropic resolution, FOV = 256 mm, TR = 3205 ms, TE_1-4_ = 1.888, 3.832, 5.776, 7.72 ms, TI = 1150 ms, FA = 7°, bandwidth = 651 Hz per pixel, and GRAPPA = 2.

### MRI Data Processing

We used the automated recon-all pipeline of FreeSurfer (version 7.1.0) to process T1w MRI scans of the participants from the HCP, SCA and IGD datasets and of FreeSurfer (version 6.0.0) to process the T1w MRI scans of the participants from the UKB dataset This pipeline generates cortical surfaces from which morphometric features (i.e., CV, SA and CT) were measured (as previously described in^74^) for 360 cortical parcels in the multimodally-derived HCP regional parcellation^36^. To control for overall brain size, we computed global CV and SA (by summation over 360 regions) and area-weighted mean CT (by averaging over 360 regions weighted by their surface area) for use in statistical models as a covariate. To control for variation in image and surface reconstruction quality, all included scans were verified as having Euler numbers (an index of topological defects in cortical surface reconstruction) less than the recommended cut off of -217^75^ and Euler number was included as a covariate in all statistical models estimating group differences in neuroanatomy (see below).

### Statistical Analyses

#### Estimating Sex Effects on Regional Cortical Anatomy

All statistical analyses were performed in MATLAB (version 2023a). We quantified the standardized effects of sex differences (females vs. males) within the HCP and UKB datasets. For the HCP dataset, a linear statistical model was defined as: *Feature ∼ Sex + Age + Euler + Global Feature*, where Feature is regional CV, SA or CT, Global Feature is global CV, global SA or area-weighted global mean CT, Age is age at scanning, and Euler is Euler number. Sex was defined as 1 for females (XX) and 0 for males (XY). All dependent variables were z-normalized across individuals so that the beta coefficient associated with sex represents a standardized effects size. For the UKB dataset, the same model with an additional term *Site* was used to account for various research sites in which the participants were scanned.

#### Defining X and Y Dosage Effects on Regional Cortical Anatomy

Case-control differences (XXY vs. XY and XYY vs. XY) quantified X and Y dosage effects (respectively) within the SCA dataset. We applied a linear model for each case-control cohort defined as: *Feature ∼ Group + Age + Euler + Global Feature*, where Group is defined as 1 for XXY or XYY and 0 for XY. All dependent variables were z-normalized across individuals so that the beta coefficient associated with Group represents a standardized effects size for a unit increase in X- and Y-chromosome dosage.

#### Defining the Effects of Intact Testicular Hormone Production on Regional Cortical Anatomy

We quantified the effects of intact testicular hormone production (typically developing XY controls vs. IGD XY individuals) using a linear model of the form: *Feature ∼ Group + Age + Euler + Global Feature*, where Group is defined as 1 for typical XY males and 0 for IGD male patients. All dependent variables were z-normalized across individuals so that the beta coefficient associated with Group represents the standardized effects size of intact testicular hormone production.

#### Controlling for Multiple Comparisons

We used FDR correction^39^ with p_FDR_ (the expected proportion of falsely rejected null hypotheses) set at 0.05 to account for multiple comparisons across cortical regions in analyses of sex, sex chromosome aneuploidy and isolated GnRH deficiency effects on CV, SA and CT. For multiple comparisons across cortical features - e.g. when spatial correlation analyses (see below) were repeated for CV, SA and CT sex difference maps - we computed a Bonferroni corrected p-value (p_BF_=3*p), which was thresholded for statistical significance at p_BF_ = 0.05.

#### Constructing Null Distributions for Comparing Two Cortical Maps

We used two complementary non-parametric approaches to test for statistical significance of observed spatial correlations between cortical maps.

The first approach used Spin Tests^37^ in which an observed spatial correlation between two cortical maps was compared to the null distribution of 10,000 spatial correlations from random spin-based spatial permutations of one of the cortical maps. This comparison yields an empirical p value (p_SPIN_), which was then further corrected across analyses for CV, SA and CT by computing a Bonferroni corrected p-value (p_SPIN-BF_=3*p_SPIN_) to be thresholded for statistical significance at p_SPIN_-_BF_ = 0.05.

The second approach relied on group permutations and could be implemented when at least one of the maps was generated by comparing two groups (e.g. a map of cortical sex differences). In this approach, the observed spatial correlation between two cortical maps was compared to the null distribution of correlations for each of 10,000 random permutations of group assignment (e.g. sex or case-control assignment in sex chromosome aneuploidy and IGD analyses) used to generate one of the maps. This comparison yielded an empirical value (p_PERM_), which was then further corrected across analyses for CV, SA and CT by computing a Bonferroni corrected p-value (p_PERM-BF_=3*p_PERM_) to be thresholded for statistical significance at p_PERM-BF_ = 0.05.

**Sup Table 2** summarizes the usage of these two non-parametric nulls across the test included in this paper.

### Quantifying the Relationships between Sex Differences in CV, SA and CT

We computed cortex-wide Pearson correlations to quantify the spatial concordance between sex differences for each unique pair of cortical features (i.e. CV vs. SA, CV vs. CT and SA vs. CT). The statistical significance of the correlations was determined by calculation of both p_PERM_ and p_SPIN_ as described above, with additional Bonferroni correction (p_SPIN-BF_ and p_PERM-BF,_ thresholded at 0.05) across the 3 inter-feature comparisons.

We then quantified the relative contributions of sex differences in SA and CT to all regions showing a statistically significant sex difference in CV (p_FDR_<0.05) by calculating the log_10_-ratio between the absolute difference between CV and SA effect sizes and the absolute difference between CV and CT effect sizes. Positive ratios indicate that CT drives sex differences in CV, whereas negative ratios indicate that SA drives sex differences in CV. We also sought to quantity the extent to which sex-biased cortical anatomy is missed by solely focusing on the predominantly used measure of CV. To that end, we performed a conjunction analysis to examine whether the regions of insignificant CV sex differences (p_FDR_>0.05) have significant (p_FDR_<0.05) sex differences in SA and CT (significant in isolation, both significant in the same direction, both significant in the opposite directions or both insignificant) and quantified the proportions of these regions that make up the regions of insignificant CV sex differences.

### Reproducibility of Sex Differences within the HCP Dataset

We quantitatively assessed the within-sample reproducibility of sex differences in CV, SA and CT using 10,000 instances of split-half, sex-balanced random subsampling of the HCP dataset at varying sample sizes (from 50 to 450, with increments of 50) and without replacement. For each instance, sex differences were recomputed in each independent split-half of the HCP dataset (using the approach described in “*Estimating Sex Effects on Regional Cortical Anatomy”* above). We then calculated the cortex-wide Pearson correlation of the standardized effects of sex differences between the split-halves (separately for CV, SA and CT). This procedure generated a distribution of 10,000 spatial correlations for each sample size, thereby offering an objective measure of the within-sample replicability of sex differences in cortical anatomy and its relation to sample size.

### Reproducibility of Sex Differences between HCP and UKB Datasets

To assess out-of-sample reproducibility of sex-biased CV, SA and CT, we computed the cortex-wide Pearson correlations between the sex differences computed in HCP and UKB datasets for each cortical feature. The statistical significance of these correlations was determined based on the null distributions derived from spin-based spatial permutations (p_SPIN_) and sex permutations (p_PERM_) of the HCP sex differences, followed by Bonferroni correction at 0.05 across the 3 cortical features (p_SPIN-BF_ and p_PERM-BF_).

We then examined the regional congruence of sex differences found between the HCP and UKB datasets. The approach was based on a conjunction analysis, for which we defined 3 levels of agreement between the two datasets (hard: UKB and HCP both showing a statistically significant sex difference in the same direction; neutral: UKB and HCP both showing the same direction of sex difference, but only reaching statistical significance in one dataset; soft: UKB and HCP both showing the same direction of statistically insignificant sex difference) and 3 levels of disagreement (hard: UKB and HCP showing a statistically significant sex difference in opposite directions; neutral: UKB and HCP showing a sex difference in opposite directions which reached statistical significance in one dataset; soft: UKB and HCP showing statistically insignificant sex differences in opposite directions). To quantify the extent to which sex-biased cortical anatomy agrees between the two datasets, we calculated the proportion of cortical regions that fell into each of these possible conjunction agreement/disagreement cases. To determine the statistical significance of regional agreement between the two datasets, we compared the combined proportion of the cortical sheet in agreement (in all 3 levels) to the null distribution of agreement proportions after 10,000 spin-permuted HCP sex difference maps, followed by Bonferroni correction across the 3 cortical features. Note that the null model based on recomputing the sex difference maps after 10,000 permutations of sex led to varying number of significantly sex-biased regions, which rendered sex permutations not applicable for determining the significance of the combined proportion of cortical sheet in agreements.

### Functional and Molecular Signatures of Sex Differences in Cortical Anatomy

#### In vivo resting-state functional networks

To assess the *in vivo* functional signatures of the sex-biased CV, SA and CT, we used a projection of the Yeo-Krienen 17 resting-state functional networks^41^ onto the same Glasser parcellation in which we have estimated regional neuroanatomical sex differences^76^. We then calculated the mean standardized effect sizes of sex differences within each Yeo-Krienen 17 resting-state functional network (i.e., somatomotor A and B, dorsal attention A and B, frontoparietal A, B and C, salience/ventral attention A and B, limbic A and B, visual A and B, default mode A, B and C, temporoparietal^41^) for the 3 cortical features: CV, SA and CT. The statistical significance of these effect sizes was determined relative to the null distribution of the effect sizes computed from the 10,000 spin-based spatial permutations of the functional networks and sex differences in CV, SA and CT (p_SPIN_), followed by Bonferroni correction (p_SPIN-BF_) at 0.05 across the 3 cortical features to control for multiple comparisons.

#### Molecular pathways related to neurodevelopment

To rank cortical expressed genes by their spatial covariation with regional neuroanatomical sex differences, we used postmortem measurements of gene expression from the AHBA (six postmortem adult brains; one female; age = 42.5 ± 13.4 years)^42^. In brief, this microarray transcriptomic dataset was acquired using 58,692 probes at 1,304 spatially distributed bulk tissue samples, which we then mapped to 20,781 unique genes from the left hemispheres of the 6 donors as described in prior work^43^. The gene expression profiles were then converted into dense expression maps as detailed by Wagstyl et al^43^. Briefly, this involved reconstructing cortical surface models on the native MRI of each donor, extracting gene expression data at nearest mid-surface cortical vertices for each bulk tissue sample (excluding samples further than 20 mm from the mid-surface vertices) and interpolating expression values for cortical vertices that weren’t directly sampled using their nearest sampled neighbor vertex. These densely sampled gene expression data was averaged across the six donors (Y-linked gene expression was extracted from male donors only) and vertex-level expression estimated were averaged for each of 360 cortical regions of the multimodal HCP parcellation^77^. We used expression maps for the set of 16,573 genes classified as having documented brain expression by Wagstyl et al.^43^ based on prior proteomic and transcriptomic studies. These preprocessing steps of the AHBA dataset enabled us to rank genes by the spatial correlation between their cortical expression and regional variation in sex effects on cortical anatomy. This ranking was done for each of 6 sets of cortical regions: female-biased and male-biased (i.e., those with positive or negative beta coefficients for the effect of sex, respectively) CV, SA and CT. We used gene set enrichment analysis (GSEA)^44^ to tests if *a priori* gene sets of interest were over-represented within each of the 6 ranked gene lists. The *a priori* gene sets of interest were drawn from a previous study and represented sets of genes tagging different epochs of brain development, cortical layers, cell types and subcellular compartments^43^. Specifically, these gene sets include: i) gene sets with peak expression in fetal (8-24 post conception week), peri-natal (24 post-conception week – 6 months post-natal) and post-natal (>6 months pos-natal) epochs of brain development^78^; ii) sets of genes having the top 5% expressions for 7 transient fetal cortical layers (subpial granular zone (SG), marginal zone (MZ), outer and inner cortical plate, subplate zone, intermediate zone, outer and inner subventricular zone, and ventricular zone)^79^; iii) marker gene sets for fetal cells (excitatory deep layer, excitatory maturing, excitatory migrating, intermediate progenitor, interneuron caudal ganglionic eminence, interneuron medial ganglionic eminence, microglia, oligodendrocyte progenitor cell, pericytes, cycling progenitor (G2- and M-phase), cycling progenitor (S-phase), outer radial glia, ventricular radial glia) from single-cell transcriptomic atlas of human cortex during mid-gestation^80^; iv) adult marker gene sets for 6 cortical layers defined as a union of two transcriptomic atlases of prefrontal cortex^81,82^; v) adult cortical cell type marker gene sets integrated from multiple single-cell transcriptomic datasets covering whole the cortex^83–90^ and fitted into excitatory neurons, inhibitory neurons, oligodendrocytes, astrocyte, oligodendrocyte precursor cells, microglia, and endothelial cells; vi) gene sets for cellular compartments (cytoskeleton, cytosol, reticulum, endosome, extracellular space, Golgi apparatus, lysosome, mitochondrion, nucleus, peroxisome, plasma membrane) were taken from the COMPARTMENTS database^91^, which integrates Human Protein Atlas, literature mining and GO annotations; vii) cellular compartment for neuronal synapse was generated by pulling all genes in the manually curated SynGO dataset^92^. For a stringent test for statistical significance, we compared the p value for each gene set to the null distribution of p values by repeating GSEA on 6 ranked gene lists after 10,000 spin-based spatial permutations of sex difference maps (p_SPIN_<0.05), followed by Bonferroni correction across 6 ranked gene lists to control for multiple comparisons (p_SPIN-BF_<0.05).

### Relationship of Sex Differences in Cortical Anatomy with X/Y Dosage and Testicular Hormone Production

We annotated regions of statistically significant sex differences in cortical anatomy by the regional congruence of X, Y and T effects (as estimated in neuroimaging analyses of SCA and IGD datasets described above). Specifically, we categorized all cortical regions according to whether the direction of sex differences was congruent with the significance and direction of X, Y and T effects. Congruence was examined as follows. For regions of female-biased anatomy, observation of anatomical increases in XXY vs. XY, anatomical decreases in XYY vs. XY, and anatomical decreases in male controls vs. males with IGD were considered congruent. Conversely, for regions of male-biased anatomy, observation of anatomical decreases in XXY vs, XY, anatomical increases in XYY vs. XY, and anatomical increases in controls vs. IGD were considered congruent. We defined two levels of congruence. Hard congruence was defined as statistically significant X, Y and T effects in congruent direction for sex differences. Soft congruence was defined as statistically insignificant X, Y and T effects in congruent direction. For each cortical feature, we computed the proportion at which each of the categories make up the significantly female-biased and male-biased cortical sheets. These proportions were then compared to the null distribution of the proportions computed after 10,000 spin-based spatial permutations of sex difference maps to determine statistical significance (p_SPIN_<0.05), followed by Bonferroni correction (p_SPIN-BF_<0.05) across the 3 cortical features to control for multiple comparisons.

## Acknowledgments

We thank all study participants for taking part in this research. This research was supported by the Intramural Research Program of the NIMH (1ZIAMH002949-09).

