## Supplementary Information for "Fractionation of sex differences in human cortical anatomy"

|  | **HCP** | | **UKB** | | **SCA XXY** | | **SCA XYY** | | **IGD** | |
| --- | --- | --- | --- | --- | --- | --- | --- | --- | --- | --- |
|  | **Female** | **Male** | **Female** | **Male** | **XXY** | **XY** | **XYY** | **XY** | **IGD XY** | **XY** |
| Sample size (n) | 592 | 493 | 375 | 294 | 99 | 92 | 34 | 47 | 19 | 22 |
| Age  (mean ± SD) | 29.56 ± 3.6 | 27.89 ± 3.59 | 48.75 ± 0.94 | 48.89 ± 0.83 | 16.38 ± 4.78 | 16.24 ± 5.64 | 15.49 ± 5.33 | 13.97 ± 4.76 | 23.27 ± 8.33 | 21.89 ± 7.66 |

**Supplementary Table 1**. Demographic characteristics of Human Connectome Project (HCP), UK Biobank (UKB), Sex Chromosome Aneuploidy (SCA) and isolated gonadotropin-releasing hormone (GnRH) deficiency (IGD) datasets.

| **Analysis** | **Statistical Tests Applied** |
| --- | --- |
| Figures 1B, 2C | Comparison of cortical maps using both Spin and participant permutation tests (P_SPIN_ and P_PERM_, respectively), with subsequent Bonferroni correction of these empirical p values (P_SPIN-BF_ and P_PERM-BF_ < 0.05) across the 3 cortical features, CV, SA and CT. |
| Figures 1C, 2D, Figure 4A, 4B, 4C (first rows) | The statistical significance of group (females vs. males, XXY vs. XY, XYY vs. XY, typical XY vs. XY IGD) effects is controlled for multiple comparisons using FDR correction (p_FDR_<0.05) across 360 cortical regions. |
| Figures 2E, Figure 4A, 4B, 4C (second rows) | Proportion of cortical sheets in agreement or congruence using Spin permutation test (p_SPIN_), with subsequent Bonferroni correction of these empirical p values (P_SPIN-BF_<0.05) across the 3 cortical features, CV, SA and CT. |
| Figure 3A | The mean standardized effect sizes of cortical sex differences within each of the 17 functional networks using Spin permutation test (p_SPIN_), with subsequent Bonferroni correction (p_SPIN-BF_<0.05) across 3 test cases (CV, SA and CT). |
| Figure 3B | Gene set enrichment using Spin permutation test (p_SPIN_), with subsequent Bonferroni correction (p_SPIN-BF_<0.05) across 6 test cases (female-bases vs. male-biases for CV, SA and CT). |
| Sup Figure 1 | Comparison of global phenotypes using linear correlation (p), with subsequent Bonferroni correction across the 3 cortical features, CV, SA and CT. |

**Supplementary Table 2.** Summary of statistical testing and multiple correction methods used in this study.

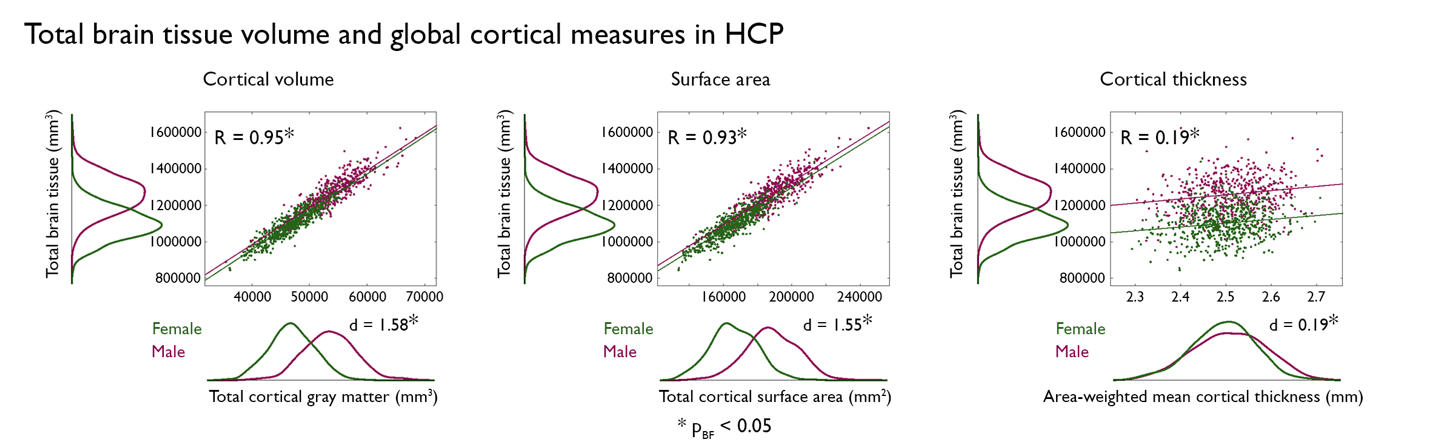

**Supplementary Figure 1. Relationships between total brain tissue volume and global surface-based measures of cortical anatomy show the sex difference in each global phenotype and the limitations of total tissue volume as a global size covariate in analysis of cortical thickness**. Scatter plots showing the relationship between interindividual variation in total brain tissue volume and global measures of cortical volume, surface area and thickness (females – green; males – pink), with least squares linear regression fit lines in each sex. Marginal histograms show the distribution of each feature in each sex, with Cohen’s effect size (d) comparing each feature between the sexes, and asterisks denoting Bonferroni-corrected statistical significance across plots at 0.05. The Pearson correlation (R) between features is shown for each plot with asterisks denoting Bonferroni-corrected statistical significance across plots at 0.05. The low R value for cortical thickness shows why total brain volume is not an equally effective covariate for control of global size in analyses of cortical volume, surface area and thickness.

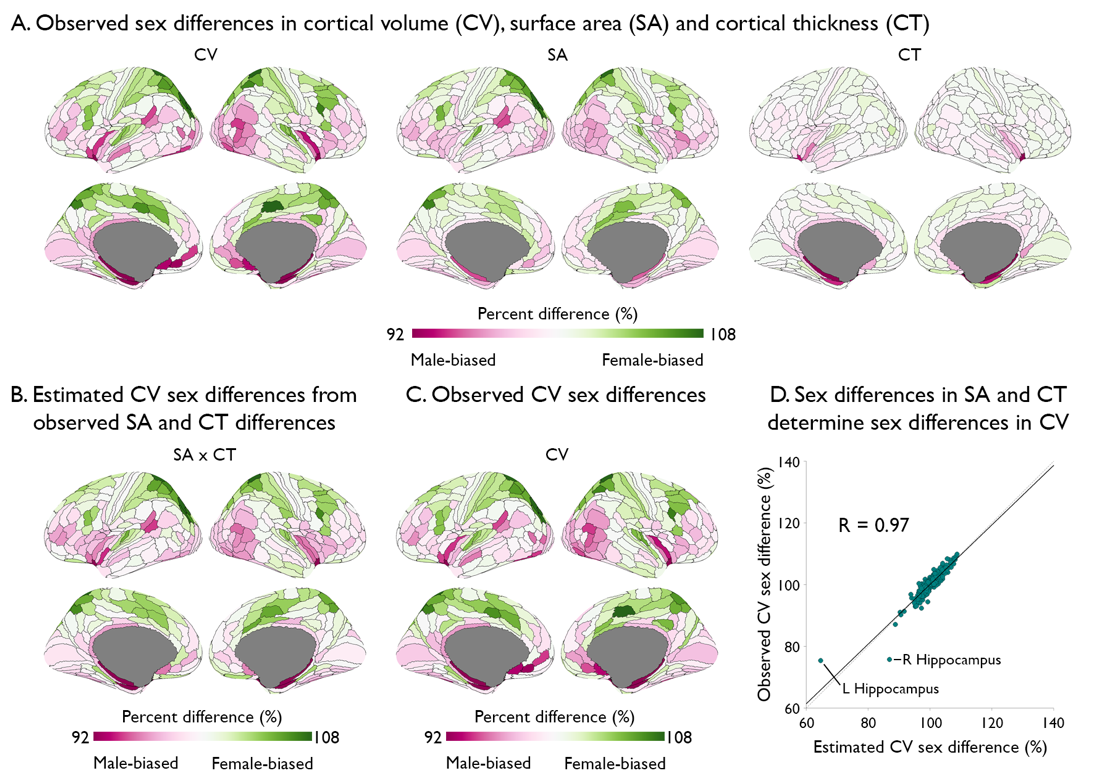

**Supplementary Figure 2. Sex differences in cortical volume (CV) are the product of sex differences in cortical surface area and thickness (SA and CT)**. **A**. Observed percent sex difference in CV, SA and CT across 360 cortical regions. **B.** The product of percentage difference maps for SA and CT. **C.** Observed percent sex difference in CV across 360 cortical regions from A. **D.** A scatter plot verifying the expected high concordance (Pearson R = 0.97) between the observed percentage sex difference in regional CV (y-axis) and the multiplication of percentage sex difference maps for SA and CT (x-axis). The solid black line is the least squares linear regression fit line – which perfectly overlays and therefore obscures the dotted line x=y identity line. Note that directly estimated sex differences in CV are least well predicted by observed sex differences in SA and CT for the left and right hippocampus.

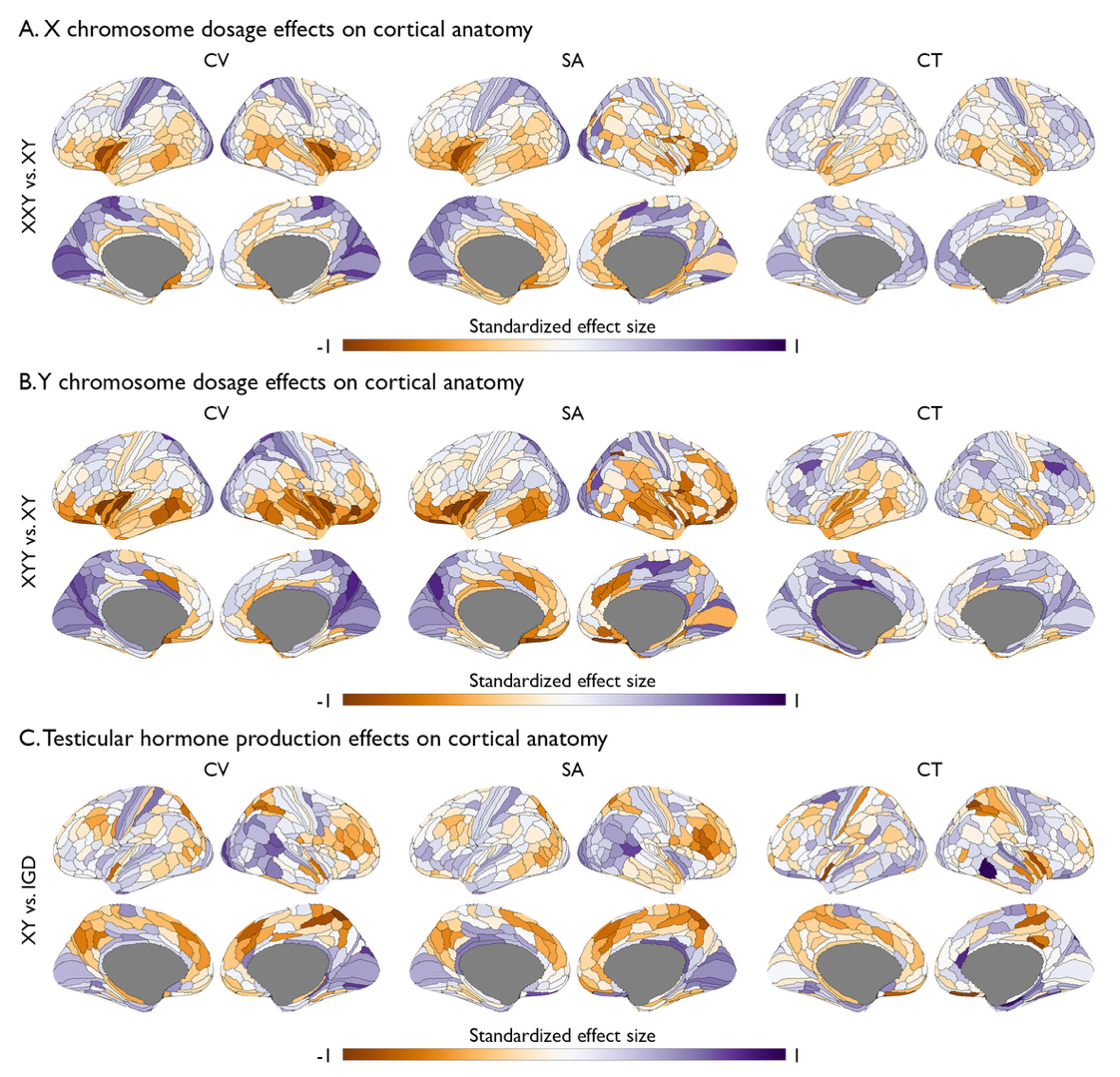

**Supplementary Figure 3. The effects of X and Y chromosome dosage and testicular hormone production on regional cortical volume (CV), area (SA) and thickness (CT)**. **A**. Standardized effect sizes across 360 cortical regions showing X-chromosome dosage (XXY vs. XY) effects on CV, SA and CT **B**. Same as panel **A** but for Y-chromosome dosage effects (XYY vs. XY). **C.** Same as panel **A** but for testicular hormone production effects (male controls vs. males with IGD).
